# Arabidopsis hydathodes contain a dense and heterogeneous epithem for apoplastic fluid release

**DOI:** 10.64898/2026.06.02.729171

**Authors:** Hiroki Yagi, Iori Mihara, Takato Ikeda, Ryosuke Sano, Taku Demura, Ikuko Hara-Nishimura, Tomoo Shimada, Haruko Ueda

## Abstract

Hydathodes are specialized leaf tissues that mediate guttation, the release of liquid water droplets from vascular plants. They consist of xylem endings, water pores, and the epithem. The epithem comprises small cells whose cell types and functional properties remain poorly understood. While molecular genetic information on hydathodes has accumulated—particularly in the model plant *Arabidopsis thaliana* through recent RNA-seq analyses—high-resolution anatomical insights into their spatial organization remain limited. In this study, we aimed to characterize the specialized anatomy of Arabidopsis hydathodes using complementary imaging approaches. A fluorescent dye taken up from roots stained hydathodes, consistent with a connection between hydathodes and the vasculature. Microfocus–X-ray computed tomography revealed that the hydathodes protrude from the abaxial leaf surface and are more densely packed internally than surrounding tissues. Light microscopy showed that the epithem was heterogeneous, with elongated cells proximally and rounded cells distally. We found that the epithem was surrounded by cells (hereafter referred to as “boundary cells”) with plastids that resembled those of mesophyll cells. Transmission electron microscopy showed that the organelles in epithem cells were immature, and the apoplast contained electron-dense material. Finally, using transgenic plants expressing a secreted fluorescent marker protein, we detected GFP in guttation droplets, indicating that the droplets can contain apoplastic material. Together, these findings reveal that Arabidopsis hydathodes possess a dense and spatially heterogeneous epithem associated with the release of apoplastic fluid.

## Introduction

Vascular plants, including *Arabidopsis thaliana*, commonly possess hydathodes at leaf tooth tips. Plants release water droplets from hydathodes, a phenomenon known as guttation. Hydathodes comprise three primary components: epithems, branched xylem endings, and water pores (hydathode pores). The epithem is composed of small, colorless, thin-walled cells (Cerutti et al. 2017; Ivanoff 1963; Jauneau et al. 2020; Singh 2014). Epithems are in direct contact with the xylem endings, and water is transported through the xylem vessels by root pressure to reach the epithems (Grunwald et al. 2003; Yagi et al. 2021a). In contrast to the xylem, the phloem is not thought to be directly connected to the epithem. This is supported by confocal imaging using transgenic Arabidopsis expressing phloem fluorescent markers (Yagi et al. 2021a). A water pore consists of a pair of guard cells similar to those of stomata. While stomata facilitate gas exchange, water pores are thought to allow liquid water to pass through. Water pores open in response to light and close in response to abscisic acid treatment (Cerutti et al. 2017). Water pores are also recognized as entry sites for bacterial pathogens (Cerutti et al. 2017; Hugouvieux et al. 1998), and they fail to close in response to flg22 in Arabidopsis (Cerutti et al. 2017).

Hydathodes have recently emerged as a subject of interest among plant researchers, particularly within the field of molecular biology. We recently revealed molecular insights into the relationship between hydathode development and auxin biosynthesis using the model plant Arabidopsis. We showed that auxin biosynthesis and accumulation precede epithem development by simultaneous visualization of three markers: *YUCCA4* promoter (auxin biosynthesis), DR5 (auxin maxima), and E325 (epithem) (Yagi et al. 2021b). Several transcriptome datasets focused on Arabidopsis hydathodes have been reported (Routaboul et al. 2024; Yagi et al. 2021a). The hydathodes were isolated and RNA-seq was carried out in these studies. Single-cell RNA-seq analyses of plant leaves also revealed hydathode cell clusters with characteristic genes, such as *PURINE PERMEASE1* (*PUP1*) (Kim et al. 2021; Liu et al. 2022; Procko et al. 2022).

In Arabidopsis, while there has been a notable increase in the genetic information available regarding hydathodes in recent years, high-resolution anatomical insights into their spatial organization remain limited in comparison to other organs. One reason for this is that hydathodes are composed of many small cells and form a thick tissue mass, rendering observation of their interior challenging in the absence of techniques such as clearing (Kawamura et al. 2010; Yagi et al. 2021a). Consequently, there is a gap between our understanding of hydathode genetics and their three-dimensional structural organization. The present study aimed to characterize the anatomy of Arabidopsis hydathodes using various imaging techniques. X-ray computed tomography (CT) scanning revealed the three-dimensional structure of hydathodes. Optical and electron microscopy analyses unveiled fine details of hydathode morphology. Additionally, we showed that guttation droplets can contain apoplastic material using transgenic plants expressing a fluorescent secretory marker protein (apoplastic green fluorescent protein [apoplastic GFP]). The findings of this study provide a foundation for future studies on the morphogenesis and physiological function of Arabidopsis hydathodes.

## Materials and Methods

### Plant materials and growth conditions

The *A. thaliana* accession Columbia-0 (Col-0) was used as a wild-type plant. The enhancer trap line E325-GFP was used as a hydathode or epithem marker line (Yagi et al. 2021a, b). Arabidopsis expressing secGFP (as “apoplastic GFP” in this study) (Uemura et al. 2012), cytosolic GFP (Mano et al. 2002), and GFP-HDEL (as “endoplasmic reticulum [ER]-retained GFP” in this study) (Hayashi et al. 2001) were used to determine whether guttation droplets contained GFP. Seeds were aseptically sown on Murashige and Skoog medium with 1% sucrose and 0.5% gellan gum (Wako, Japan). Plants were grown on plates at 22°C under continuous light conditions. For observation of guttation phenomena and collection of guttation droplets, 2- or 3-week-old plants were transplanted to vermiculite (G20; NITTAI Co., Japan) supplied with a 1/2000 dilution of HYPONeX solution (HYPONeX JAPAN Ltd., Japan). Plants were kept under high humidity (∼90%) within plastic boxes.

### Microfocus–X-ray computed tomography (CT) scanning

The 6th or 7th true leaves of 3-week-old Arabidopsis plants were cut at their petioles. To prevent the leaves from drying, the leaves were placed in 0.2 mL tubes and microfocus–X-ray CT scanned using inspeXio SMX-100CT (Shimadzu, Japan) as previously reported (Kunieda et al. 2024). CT scans of approximately 4×3 mm^2^ were taken from the basal region of the leaves, including the two hydathodes. The 3D data were analyzed with VGSTUDIO and myVGL software (Hexagon, Sweden).

### Root dip staining

E325-GFP plants (2- or 3-week-old) grown on solid medium were gently lifted with tweezers, and their roots were dipped into water containing 1% (w/v) Evans blue (Nacalai Tesque, Japan) for over 8 hours (Fig. S1). The dipped samples were kept at high humidity inside a sealed box to prevent them from drying out and shrinking. The bright and fluorescent images were taken using a fluorescence macro zoom microscope (MVX10; Olympus, Japan) and a camera (Axiocam506 color; Carl Zeiss, Germany). The images were processed by ImageJ/Fiji v1.53f51/Java 1.8.0 software (Schindelin et al. 2012; National Institute of Health).

### TEM analysis

Hydathodes located at the basal side (near the petiole) of the 6th or 7th true leaves of 3-week-old E325-GFP were isolated using razors under a fluorescence stereomicroscope (SteREO Lumar V12; Carl Zeiss, Germany). Samples were fixed with 0.05 M cacodylate buffer containing 2% paraformaldehyde and 2% glutaraldehyde at 4°C overnight. After fixation, the samples were washed three times with 0.05 M cacodylate buffer for 30 min each and postfixed with 0.05 M cacodylate buffer containing 2% OsO_4_at 4°C for 3 h. The samples were dehydrated in graded ethanol solutions (50%, 70%, 90%, and 100%), infiltrated with propylene oxide (PO) twice for 30 min each, and immersed in a 7:3 mixture of PO and resin (Quetol-651; Nisshin EM, Japan) for 1 h. The tube cap was left to allow PO to volatilize. The samples were transferred to fresh 100% resin and were polymerized at 60°C for 48 h. Ultra-thin sections (80 nm) were examined using a transmission electron microscope (TEM) (JEM-1400Plus; JEOL, Japan) at 100 kV, and digital images were taken with a CCD camera (EM-14830RUBY2; JEOL, Japan).

### SEM analysis

The 6th or 7th true leaves of 3-week-old Arabidopsis leaves were treated with a NanoSuit® Elution Type I (NanoSuit Inc., Japan) and then observed by a scanning electron microscope (SEM) (JCM7000; JEOL, Japan).

### Observation of GFP fluorescence of guttation droplets

Guttation droplets on Arabidopsis leaves were collected with micropipettes and gathered in protein low binding tubes (PRokEEP; WATSON, Japan). The bright field and GFP fluorescence were observed under a macro zoom microscope (MVX10; Olympus, Japan), and images were taken using a microscope camera (Axiocam506 color; Carl Zeiss, Germany).

### SDS–PAGE and immunoblot

Guttation droplets on Arabidopsis leaves were collected by micropipettes and gathered in collection tubes. Guttation droplets (8 µL) were mixed with 2 µL of SDS–PAGE sample buffer (250 mM Tris-HCl, pH 6.8, 10% [w/v] SDS, 25% [v/v] 2-mercaptoethanol, and 50% [v/v] glycerol) and subjected to SDS–PAGE followed by immunoblot analysis as described previously (Shimada et al. 2003). Antibodies used were anti-GFP (JL8; Clontech, USA) and anti-mouse IgG HRP (GE Healthcare, USA).

## Results and Discussion

### Arabidopsis hydathodes are vascular-connected and abaxially protruding structures

Leaf teeth represent terminal points of the vasculature, and hydathodes are thought to be connected to xylem endings. Previously, a petiole dip method—in which the cut end of a petiole is immersed in a calcofluor white solution—was used to visualize the vasculature and hydathodes (Shatil-Cohen et al. 2011). Here, we developed a root dip method (Fig. S1), in which roots were immersed in a 1% (w/v) Evans blue, a largely non-permeating dye that appears dark blue under bright-field illumination and exhibits red fluorescence (excitation/emission: 647/680 nm). Although Evans blue is widely used as an indicator of dead cells in plants (Ikegawa et al. 1998, 2000), in our assay it was used as a tracer to monitor long-distance water transport through the xylem. In bright-field images, leaf veins were stained blue and the dye consistently accumulated at leaf tooth tips (Fig. 1**a**). In fluorescence images, the vasculature was clearly visualized, whereas red fluorescence from Evans blue was detected at the site marked by the epithem reporter E325-GFP, albeit less clearly than in bright-field images (Fig. 1**a**). These observations support the notion that Arabidopsis hydathodes are connected to the vasculature and receive materials transported from roots via the xylem.

**Fig. 1.**
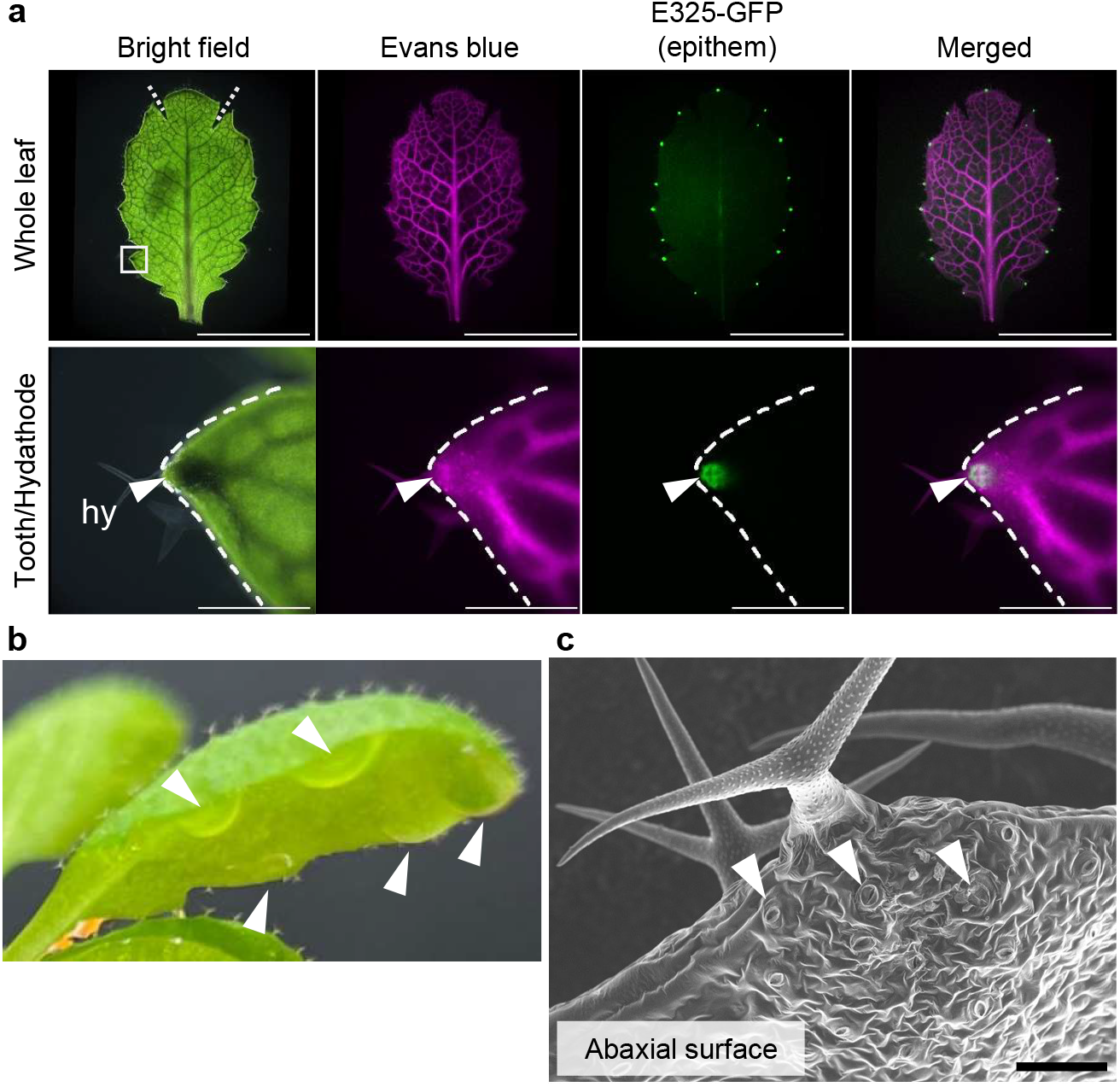
Hydathodes mediate guttation in *Arabidopsis thaliana*. **a** Root-dip staining of true leaves from E325-GFP plants (epithem marker). Upper panels: whole-leaf images including bright field (Evans blue staining), red fluorescence of Evans blue, green fluorescence of E325-GFP, and merged images; dashed lines indicate incisions made to flatten leaves. The white box in the bright-field image indicates the leaf tooth region magnified below. Lower panels: magnified tooth/hydathode region; dashed lines outline the leaf tooth. hy, hydathode. **b** Plant showing guttation droplets when grown on vermiculite under high humidity (∼90%) in a plastic box. Arrowheads indicate guttation droplets on the abaxial leaf surface. **c** SEM image of a leaf tooth viewed from the abaxial side. Arrowheads indicate water pores. Guard-cell morphology alone does not allow unambiguous discrimination between stomata and water pores. Scale bars indicate 5 mm (upper panels in **a**), 500 µm (lower panels in **a**), and 50 µm (**c**).

Arabidopsis plants produced guttation droplets on the abaxial side of leaf teeth (Fig. 1**b**). SEM analysis of the abaxial tooth surface revealed pores morphologically consistent with water pores (Fig. 1**c**), similar to previously reported SEM observations (Paauw et al. 2023; Wang et al. 2011). These pores were located near the distal region of the tooth and often appeared more open than typical stomata (Cerutti et al. 2017). However, based on surface morphology alone, it remains difficult to unambiguously distinguish water pores from stomata in Arabidopsis. Notably, guttation droplets can also emerge from the adaxial leaf surface in some eudicots. For example, Singh presented photographs of three terrestrial eudicot plants—strawberry (*Fragaria ananassa*), tomato (*Solanum lycopersicum*), and garden burnet (*Sanguisorba minor*)—in the book “Guttation” (Singh 2020), all of which exhibited droplets on the adaxial surface of the leaves. Thus, the leaf surface from which guttation droplets are released is not necessarily conserved across species.

To examine hydathode morphology non-destructively, we performed microfocus X-ray CT scanning of Arabidopsis leaves (Fig. 2, Supplementary videos 1, 2). Three-dimensional reconstructions showed that tooth tips containing hydathodes protruded toward the abaxial side (Fig. 2**a, b**). Virtual sections further indicated that the interior region of hydathodes was more densely packed than surrounding mesophyll tissues (Fig. 2**c**, S2). The dense domain was approximately 100 µm in size, which is comparable to the region labeled by E325-GFP (Yagi et al. 2021a). Together, these imaging data indicate that Arabidopsis hydathodes are dense and abaxially protruding structures.

**Fig. 2.**
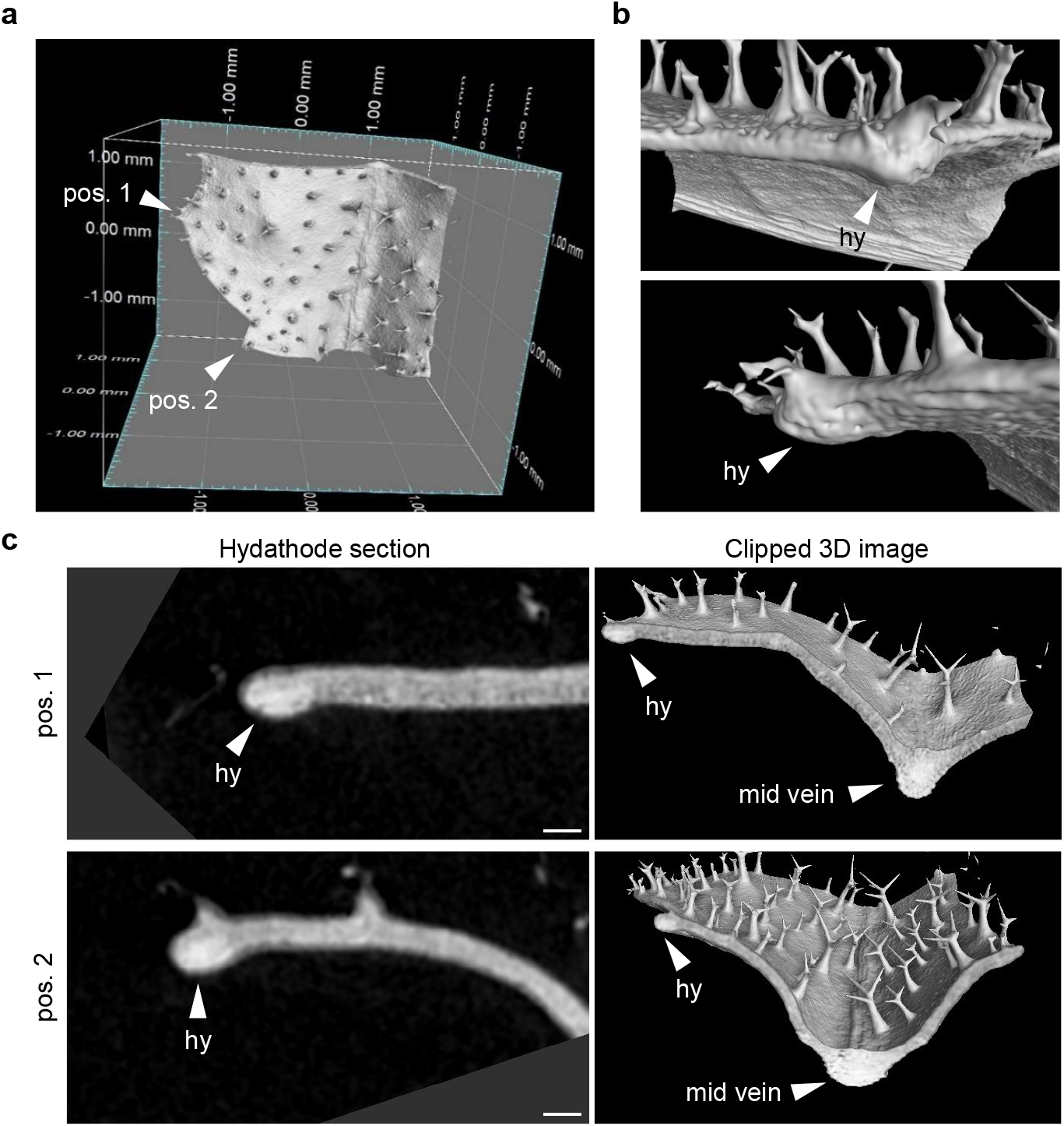
Hydathodes protrude and they are densely packed. **a** Three-dimensional image of Arabidopsis hydathodes obtained by microfocus-X-ray computed tomography (CT). 3D rendering of the basal leaf region (∼4×3 mm^2^) containing two hydathodes; arrowheads indicate hydathode at positions (pos.) 1 and 2. **b** Close-up view of the surface of the hydathode pos. 1. hy, hydathode. **c** Virtual sections (pseudo-sections) through the hydathodes. Scale bars indicate 100 µm (**c**).

### Arabidopsis epithems are closely packed and heterogeneous

In a previous study, we reported Arabidopsis hydathode structures using fluorescent protein markers (Yagi et al. 2021a). However, this approach did not allow for detailed visualization of fine anatomical features, such as apoplastic spaces and subcellular organization. To address this limitation, we conducted light microscopy of resin sections stained with toluidine blue and performed TEM analysis on Arabidopsis hydathodes (Figs. 3, 4). In vertical sections, we observed small cells in the epithem, designated as epithem cells (Fig. 3, S3. Pseudo-colored as green in Fig. 3**b**). The intercellular space between epithem cells was so narrow that it was barely discernible under light microscopy. In contrast, the intercellular space in the spongy or palisade mesophyll was clearly visible (pseudo-colored as bright red in Fig. 3**b**). Our observations revealed that epithems are composed of cells with heterogeneous morphology. The epithem cells at the distal side of the hydathodes were smaller than mesophyll cells and rounded in shape (Fig. 3**d, f**, S3). Conversely, epithem cells at the proximal side of the hydathodes (i.e., in proximity to the vasculature connection site) were elongated and even smaller than epithem cells at the distal side of the hydathodes (Fig. 3**e**, S3). TEM analysis revealed that the organelles of the epithem cells, including vacuoles and plastids, were smaller than those of the mesophyll cells (Fig. 3**g, h**). Notably, we observed larger cells surrounding the epithem (pseudo-colored as yellow in Fig. 3**b**). These cells appeared to be mesophyll cells based on their size and organelles (Fig. 3**c, g**). However, unlike typical mesophyll cells, they had very narrow intercellular spaces and adhered closely to each other, forming tight contacts with the epithem (Fig. 3, S3). These cells may act as ‘boundary cells,’ surrounding the epithem and creating a distinct region that separates it from adjacent mesophyll tissues.

**Fig. 3.**
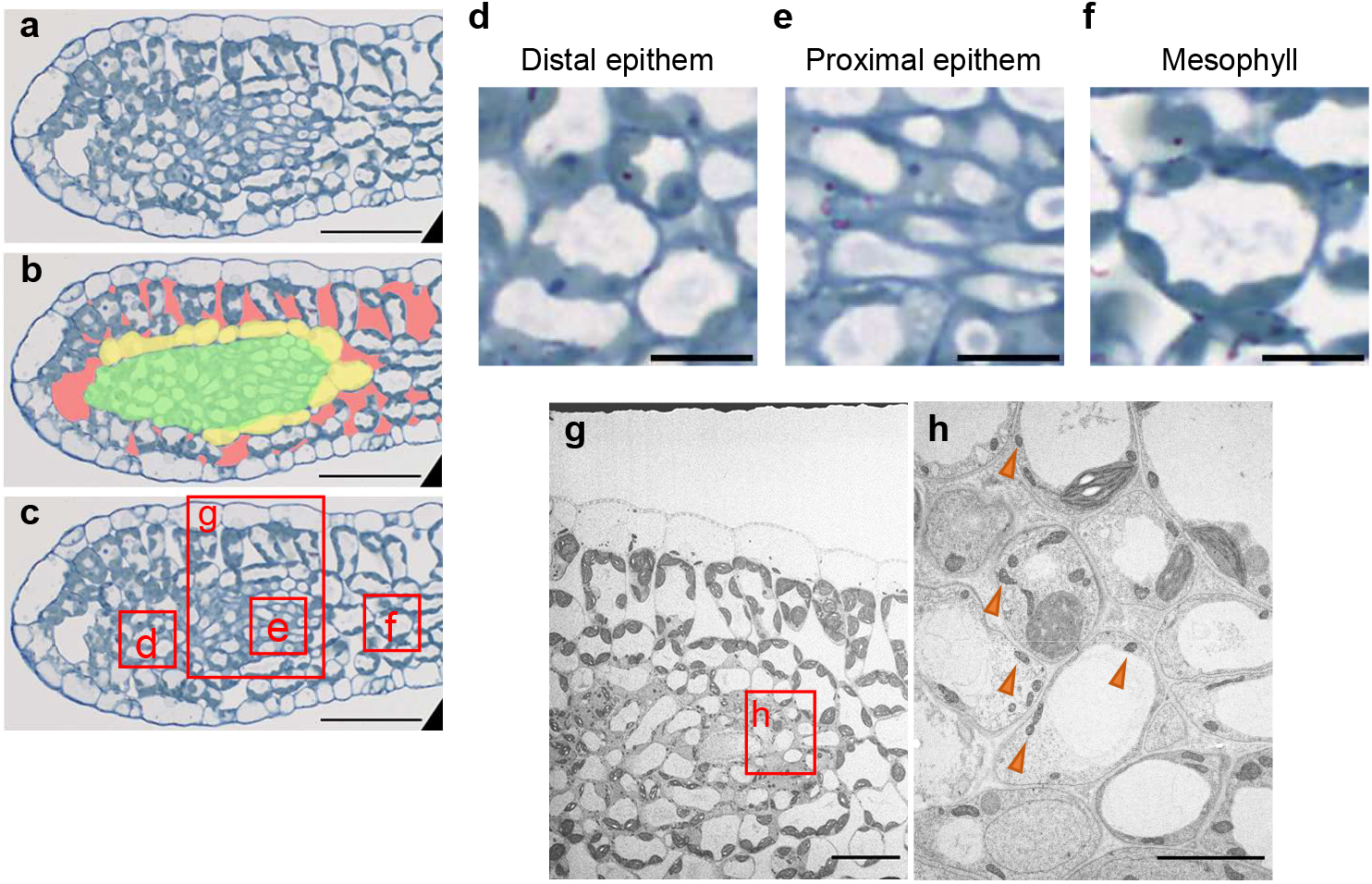
Cell-type heterogeneity and boundary structure in vertical sections of Arabidopsis hydathode. **a** A resin section stained with toluidine blue. **b** Pseudo-colored image of (**a**). Green, epithem; yellow, boundary cells; red, intercellular spaces. **c** Copy of (**a**) with boxes indicating the regions shown in (**d**–**g**). **d**–**f** Magnified views of the boxed region in (**c**). **g** TEM image of the boxed region in (**c**). **h** Higher-magnification TEM image of the boxed region in (**g**). Orange arrowheads indicate immature plastids. Scale bars indicate 100 µm (**a**–**c**), 10 µm (**d**–**f**), 20 µm (**g**), and 5 µm (**h**).

**Fig. 4.**
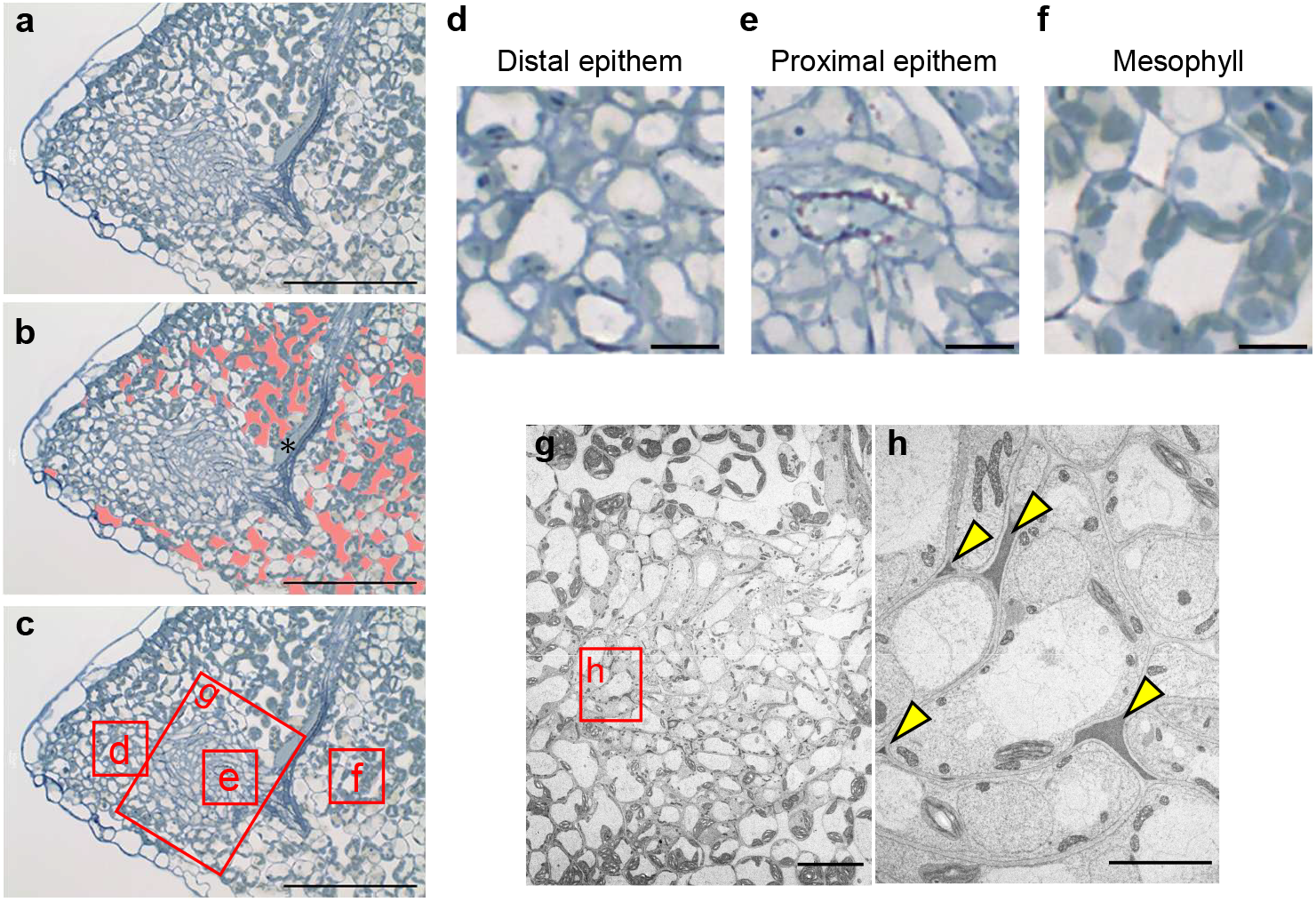
Densely packed structure and vascular penetration in horizontal sections of Arabidopsis hydathode. **a** A resin section stained with toluidine blue. **b** Pseudo-colored image of (**a**). Red, intercellular spaces. Asterisk (*) indicates a myrosin cell. **c** Copy of (**a**) with boxes indicating the regions shown in (**d**–**g**). **d**–**f** Magnified views of the boxed region in (**c**). **g** TEM image of the boxed region in (**c**). **h** Higher-magnification TEM image of the boxed region in (**g**). Yellow arrowheads indicate electron-dense material in the intercellular space. Scale bars indicate 100 µm (**a**–**c**), 10 µm (**d**–**f**), 20 µm (**g**), and 5 µm (**h**).

In horizontal sections, we observed small and closely packed epithem cells (Fig. 4, S4). Vascular strands were found to penetrate the epithem (Fig. 4**a**), a finding consistent with our previous research using confocal laser microscopy (Yagi et al. 2021a). Regions lacking intercellular spaces were clearly observed, even though boundary cells were less distinct than in vertical sections, likely due to the cutting orientation. As noted earlier, epithem cells at the distal side of the hydathodes exhibited a rounded shape, while those at the proximal side displayed an elongated shape (Fig. 4**d**–**f**, S4). The elongated cells may correspond to, or closely associate with, vascular elements since xylem endings penetrate the epithems (Cerutti et al. 2017; Yagi et al. 2021a). TEM analysis revealed that epithem cells were irregularly shaped, and the intercellular spaces between epithem cells occasionally contained electron-dense material (Fig. 4**h**). Similar high-electron-density material has been reported in laminar hydathodes of *Ficus formosana* (Chen and Chen 2006). The presence of these materials may reflect a shared structural feature of hydathodes across diverse species.

While the molecular mechanisms of leaf tooth development have been extensively studied (Bilsborough et al. 2011; Tameshige et al. 2016), the processes of cell division and differentiation within the tooth interior remain largely unexplored. The heterogeneous structure of epithems, comprising elongated cells at the proximal sides and rounded cells at the distal sides, may reflect their developmental process. Elongated cells appear sequentially along the vasculature, raising the possibility that epithem cells might derive from vascular cells rather than mesophyll cells. However, recent single-cell RNA-seq analyses do not support this hypothesis (Kim et al. 2021; Liu et al. 2022; Procko et al. 2022). These studies may have been unable to capture elongated epithem cells due to their reliance on genes such as *PUP1* or *EP3* as hydathode markers. Single-cell analyses specifically focused on hydathodes would provide new molecular insights into epithem identity. In any case, a redefinition of epithem cells based on careful observation of both morphology and gene expression is important to advance our understanding of these structures.

### Guttation droplets are derived from apoplastic fluid

Previous studies have shown that guttation droplets from barley contain proteins such as peroxidase and chitinase (Grunwald et al. 2003). These proteins feature N-terminal signal peptides, enabling their secretion into the extracellular space, or apoplast. These observations suggest that guttation droplets originate primarily from apoplastic fluid. To test this hypothesis, we employed three types of GFP constructs: (i) apoplastic GFP carrying an N-terminal signal peptide, (ii) cytosolic GFP lacking localization signals and therefore distributing throughout the cytosol and nucleus, and (iii) ER-retained GFP carrying an N-terminal signal peptide and a C-terminal HDEL motif. Our experiments revealed that guttation droplets from transgenic plants expressing apoplastic GFP displayed detectable GFP fluorescence (Fig. 5**a**). In contrast, guttation droplets from wild-type Col-0 plants and those expressing cytosolic or ER-retained GFP showed no fluorescence (Fig. 5**a**). Furthermore, immunoblot analysis using anti-GFP antibodies detected apoplastic GFP, but not the other GFP forms, in guttation droplets (Fig. 5**b**). Together, these results support the conclusion that guttation droplets are derived predominantly from apoplastic fluid rather than symplastic fluid.

**Fig. 5.**
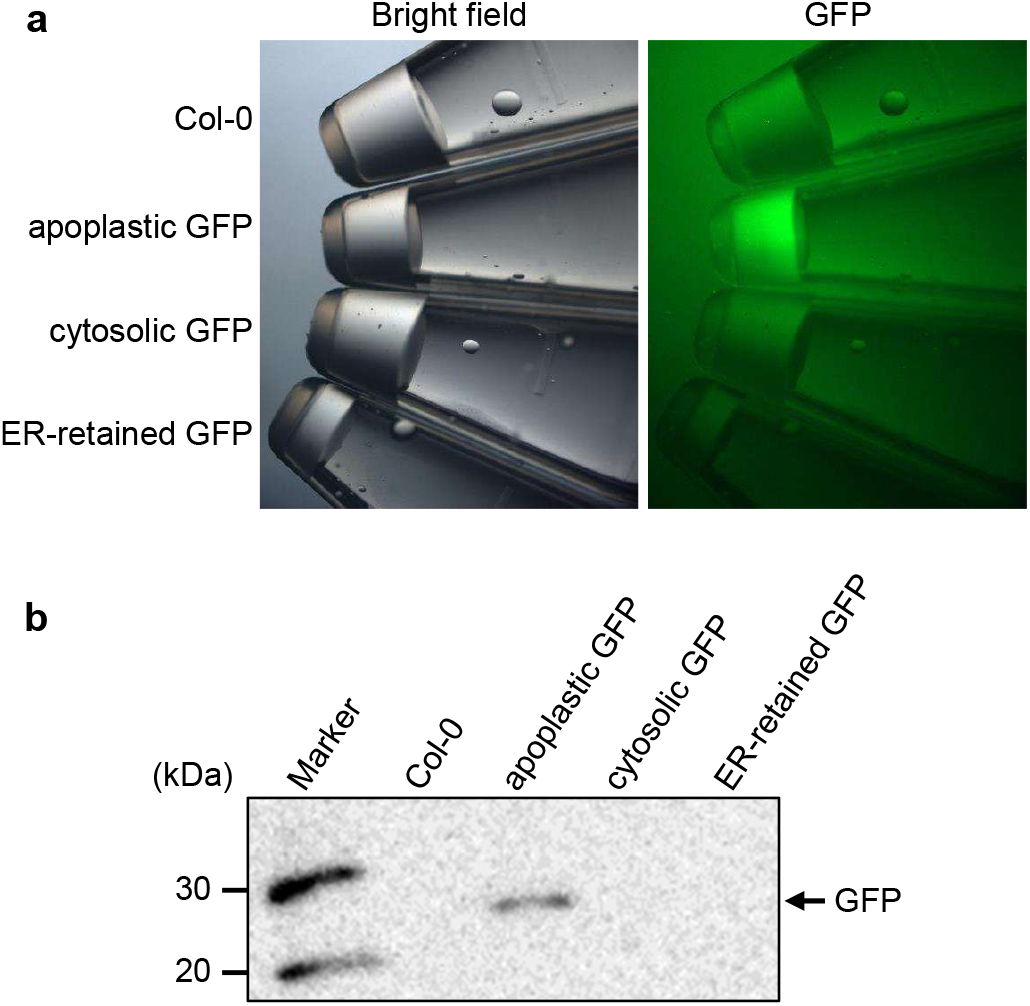
Guttation droplets are derived from apoplastic fluid. **a** Bright-field and fluorescent images of guttation droplets collected from wild-type (Col-0) plants and transgenic lines expressing apoplastic GFP, cytosolic GFP, or endoplasmic reticulum (ER)-retained GFP. **b** Immunoblot of guttation droplets showing detection of the GFP derivatives using anti-GFP antibodies.

Figure 6 illustrates our proposed structural model of the Arabidopsis hydathode. Xylem fluid flows into the epithem region and passes through narrow extracellular (apoplastic) spaces, where it can mix with apoplastic solutes. The fluid moves toward the water pores and is released to the external environment as guttation droplets. Hydathodes have been proposed to limit solute loss by retrieving valuable solutes through the action of specific transporters, including PUP1 for cytokinin, NITRATE TRANSPORTER 2.1 for nitrate, and PHOSPHATE TRANSPORTER 1;4 for inorganic phosphate (Bürkle et al. 2003; Routaboul et al. 2024). In this framework, the densely packed epithem—composed of small cells and narrow apoplastic spaces—may provide an extended interface for solute exchange and efficient retrieval before exudation occurs.

**Fig. 6.**
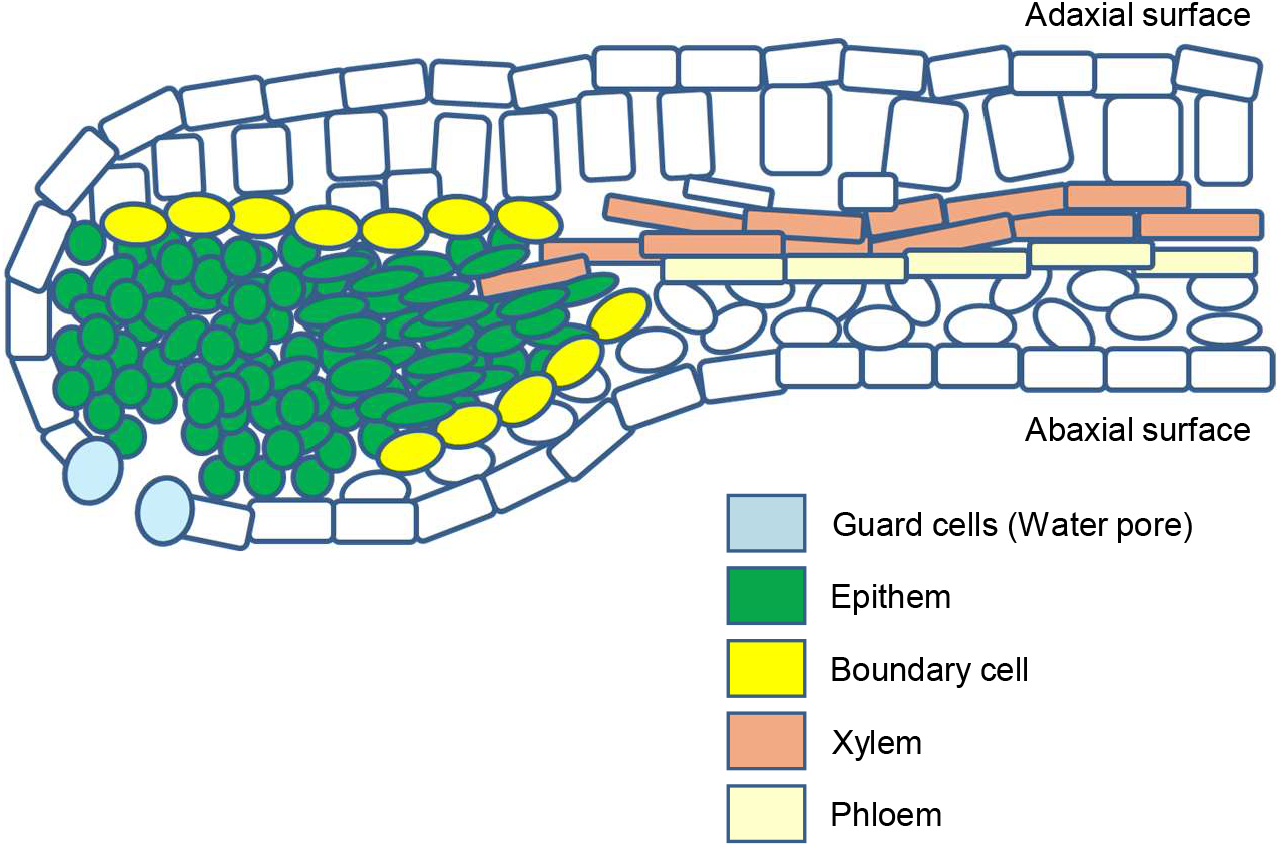
Schematic model of Arabidopsis hydathode anatomy. Hydathodes protrude on the abaxial side of the leaf. Xylem vessels penetrate the epithem, which consists of small, tightly packed cells with narrow intercellular spaces. Proximal epithem cells near the vasculature are elongated, whereas distal epithem cells near the water pores are rounded. The epithem is surrounded by boundary cells, which may prevent fluid leakage into adjacent mesophyll tissues.

Boundary cells located adjacent to the epithem may serve as a barrier, preventing fluid leakage into surrounding mesophyll tissues. This idea is consistent with both root dip staining (Fig. 1**a**) and previous petiole dip staining (Shatil-Cohen et al. 2011), which showed no dye leakage from hydathodes under these conditions. Interestingly, when bacteria were introduced via water pores, their colonization was limited within hydathodes, especially during early infection stages (Cerutti et al. 2017; Hugouvieux et al. 1998; Paauw et al. 2023). These observations are consistent with the presence of physical and/or apoplastic barriers at or around hydathodes. Further molecular characterization of these barrier structures is necessary to fully understand their composition and function.

## Acknowledgments

We are grateful to the Nottingham Arabidopsis Stock Centre (NASC) for E325 seeds (stock name: N70027). We thank Tomohiro Uemura (Ochanomizu Univ.) and Shoji Mano (National Institute for Basic Biology) for sharing Arabidopsis seeds. We also thank Daiske Honda (Konan Univ.) for advice on using SEM.

## Supplementary Information

**Fig. S1** Root-dip staining assay used to visualize the connection between hydathodes and the vasculature

**Fig. S2** Additional 3D microfocus X-ray CT scan images of Arabidopsis hydathodes

**Fig. S3** Additional images of toluidine blue-stained vertical sections of Arabidopsis hydathode

**Fig. S4** Additional images of toluidine blue-stained horizontal sections of Arabidopsis hydathode

**Supplementary video 1** 3D CT-scan movie of Arabidopsis hydathode

**Supplementary video 2** Pseudo-crosssection movie of Arabidopsis hydathode

## Author contributions

HY, HU, and TS conceptualized the present study. HY, TI, IM, and RS performed the experiments. TD, IH-N, HU, and TS supervised experiments. HY, HU, and TS wrote the original draft, and all authors reviewed the manuscript.

