## Supplementary Figs for "Arabidopsis hydathodes contain a dense and heterogeneous epithem for apoplastic fluid release"

**Title:**

### Fig. S1

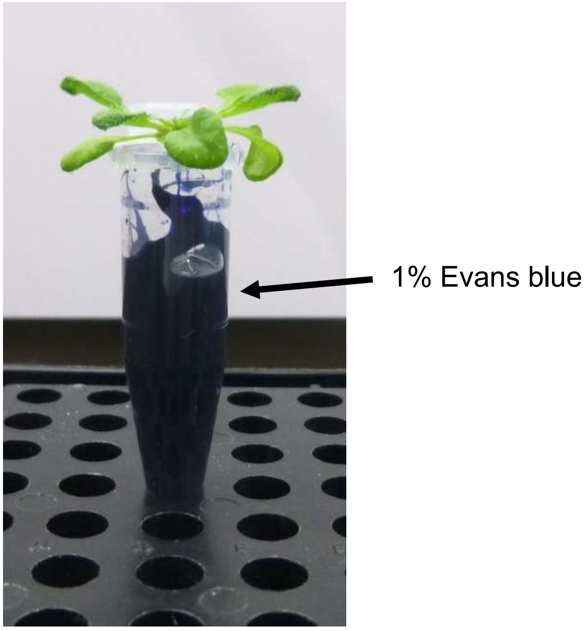


**Fig. S1** Root-dip staining assay used to visualize the connection between hydathodes and the vasculature

The image of root dip staining of Arabidopsis. Arabidopsis plants (2- or 3-week-old) grown on solid Murashige and Skoog medium were gently lifted with tweezers, and their roots were dipped into 1% (w/v) Evans blue dye solution for over 8 hours under high humidity.

### Fig. S2

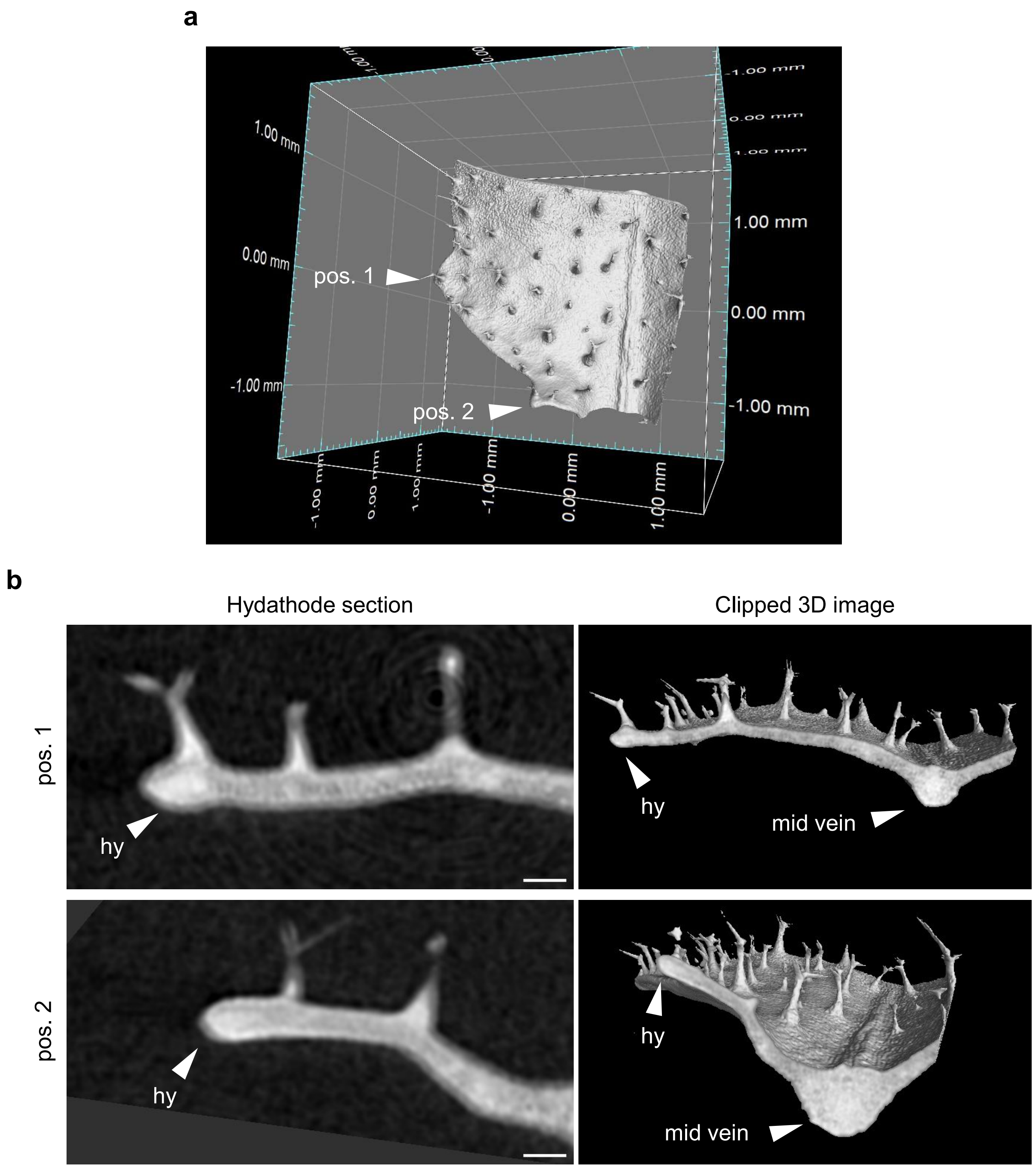


**Fig. S2** Additional 3D microfocus X-ray CT scan images of Arabidopsis hydathodes
**a** 3D rendering of the basal leaf region (~4 × 3 $mm^2$) containing two hydathodes; arrowheads indicate hydathode at positions (pos.) 1 and 2. **b** Virtual sections (pseudo-sections) through the hydathodes. Scale bars indicate 100 μm.

**Fig. S3**

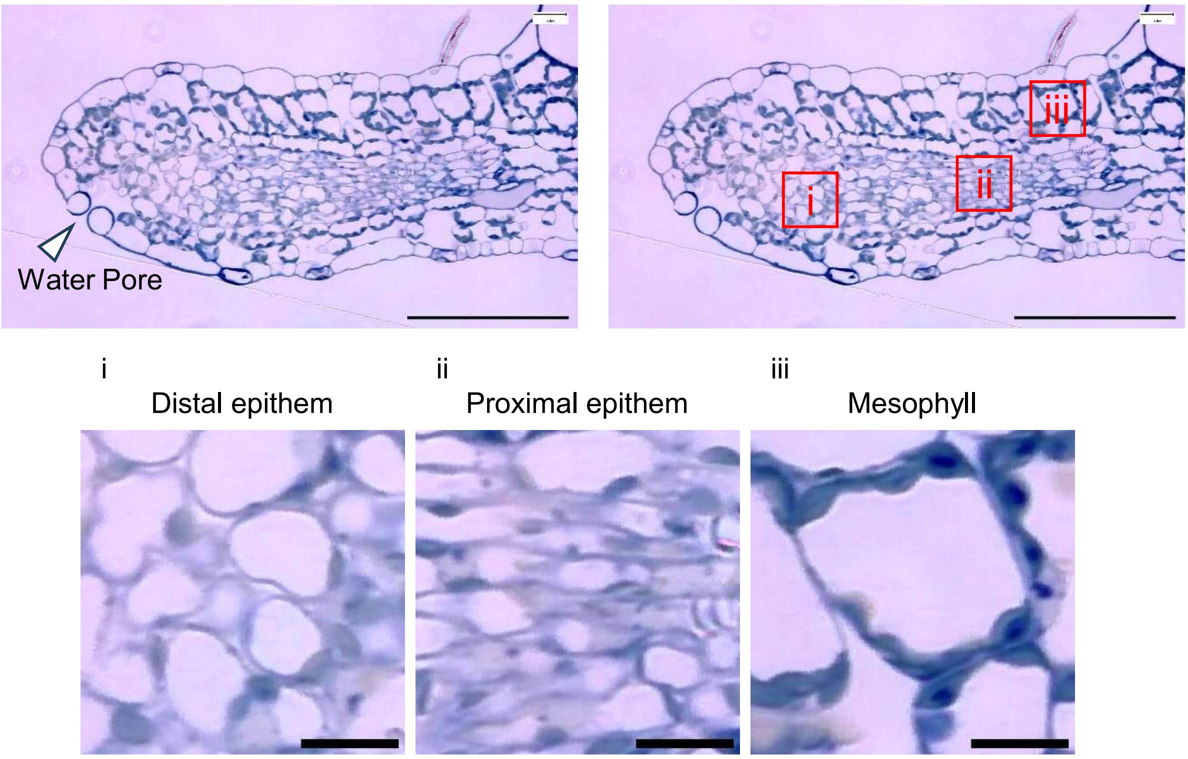


**Fig. S3** Additional images of toluidine blue-stained vertical sections of Arabidopsis hydathode
Red squares in upper right panels were magnified below. Scale bars indicate 100 µm (upper panels in each) and 10 µm (lower panels in each).

**Fig. S4**

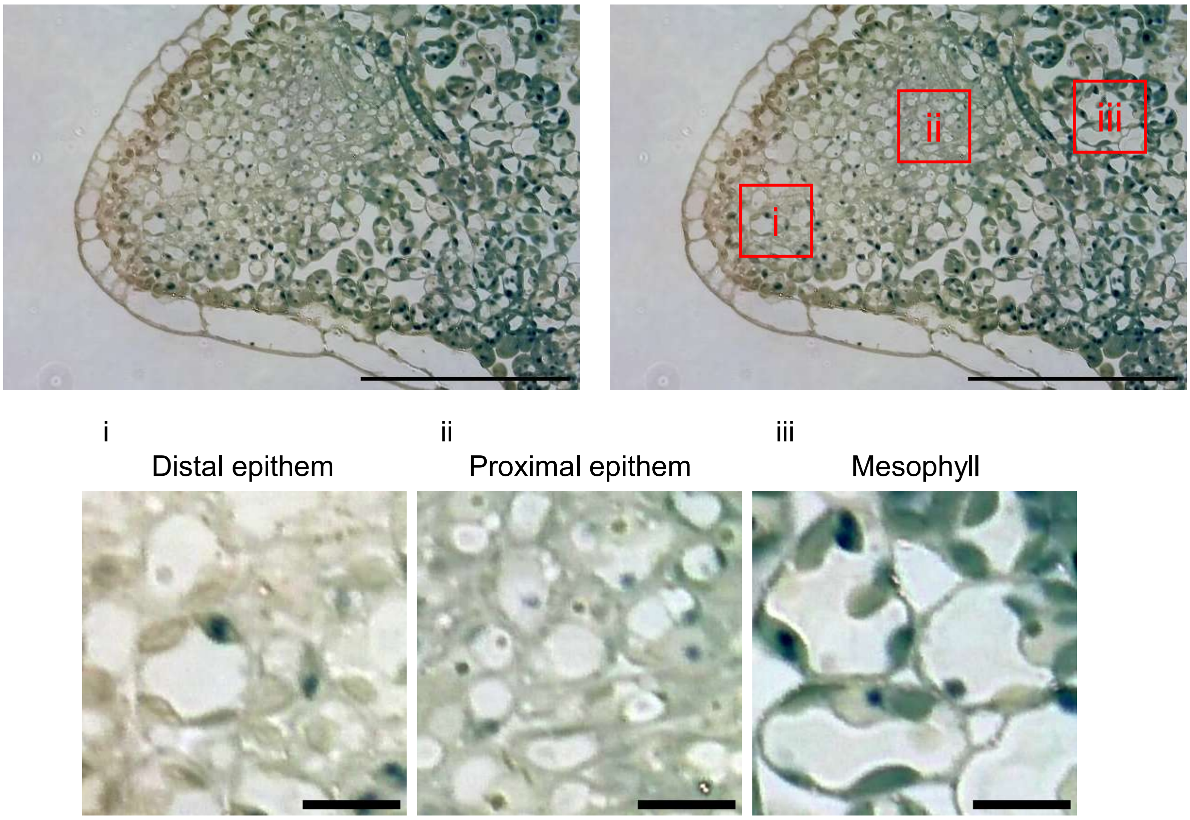


**Fig. S4** Additional images of toluidine blue-stained horizontal sections of Arabidopsis hydathode Red squares in upper right panels were magnified below. Scale bars indicate 100 µm (upper panels in each) and 10 µm (lower panels in each).
